# t-SNE transformation: a normalization method for local features of single-cell RNA-seq data

**DOI:** 10.1101/799288

**Authors:** An Chengrui, Jaume Bacardit, Zhang Nijia, Wu Bingbing

## Abstract

Single-cell RNA sequencing has been widely used by biology researchers. There are many analysis tools developed accordingly. However, almost all of them use log transformation in the process of normalization, which may result in system bias on global features of datasets. It is considered that they may not be suitable for researchers who expect local and detailed features of datasets, such as rare cell population and independent expressed genes. In this study, we developed a method called t-SNE transformation to replace log transformation. We found that it can well respond to some specific bio-markers in real datasets. When the cluster number was changed, t-SNE transformation was steadier than log transformation. Further study showed that clustering after t-SNE transformation detected the residual cells more accurately after majority cells of one type were removed manually. It was also sensitive to a highly-variated independent gene added artificially. In conclusion, t-SNE transformation is an alternative normalization for detecting local features, especially interests arouse in cell types with rare populations or highly-variated but independently expressed genes.

## Introduction

Single-cell RNA-sequencing (scRNA-seq) has been developed since 2009 [21]. It has become one of the most potent RNA-sequencing techniques to describe transcriptomic information stored in every cell of samples. It is often used to analyze global features including general cell types, their relationships within [1] and structural or functional alterations during certain biological processes [4, 25]. It can be operated by various analyzing tools such as Seurat [18], SC3 [13] or Monocle [22].

Most of the tools for analyzing scRNA-seq data use log transformation in normalization [5]. It underestimated the means of low expressed genes [15] as well as their derivations [7, 10]. The outcomes of log-transformation are the clustering results that are more tended to high expressed genes [7]. It covered some information determined by low expressed genes. Considering the high sparsity of scRNA-seq datasets, low expressed genes may be caused by their high Gini index [11]. As a result, log transformation decreases the power of clustering to detect them.

In addition, the functions of cells and genes vary greatly in different biological processes, so analyzing local features of datasets becomes essential in certain biology studies as well. In the high-dimensional space by genes, local features of scRNA-seq data can refer to subtle cell types related to the specific biological process [6]. For instance, cells with high stemness in adults’ intestine account for only less than one percent [6]. Another example is the rare existence of chemotherapy-resistant cancer cells in tumor tissues [3]. In sample space, when the expression matrix is transposed, local features become the pattern that is made up of a limited number of genes essential for certain biological processes. For example, expression of Ascl1 can inhibit neural stem cells from differentiating into retinal ganglion cell [2]. Pdx1 and Sox9 can determine whether the gut stem cells differentiate into epithelia in different parts of pancreas while removing one of them can cause the alternation of morphology [20]. What is more, by transfecting one or few transcription factors, researchers can reprogram cells from one type to another, such as Mesp1, Tbx5, Gata4, Nkx2.5 and Baf60c, transferring fibroblasts into cardiac progenitor cells [14]. Theoretically, it is the expression of these genes that can respond to environmental changes. So their correlation with other genes should be pretty low. However, the population structure of cells in tissues tested by scRNA-seq is often complex. Numerous genes express differently, generate strong multi-collinearity, of which the variation outweigh the rare but essential genes. If their correlation is low enough, such relevant genes are hard to be detected.

In this study, in order to satisfy the interests of researchers in local features of scRNA-seq data, we attempt to increase the sensitivity of scRNA-seq data analysis. The t-distributed stochastic neighbor embedding (t-SNE) is a candidate way to solve the problems considering its three functions: manifold learning, multiple Gaussian kernels and reducing dimension [16]. Manifold creates new spaces that contain the relationship within cells. It induces normalization to aim at the direct distance between cells rather than scaling gene expressions. So it avoids the occurrence of overestimating semantic information of features. Multiple Gaussian kernel methods can solve the problem of clustering heterogeneous density data [12, 23] by introducing various *θ* to fit different clusters. It also maintains the local relationship when reducing dimension. And it also keeps the local relationship when reducing dimension [23].Although t-SNE is considered as a dimension reduction method, the dimension of outcomes is not restricted to be lower than that of original space. Therefore, t-SNE transformation is defined to refer mapping data to space with the same dimension of the original space by the rule of t-SNE.

## Results

### Consequences of clustering after t-SNE or log transformation

In order to display the consequences of t-SNE or log transformation, we used three real datasets in GEO database. They are GSE81682 related to mice hematopoiesis conducted by Nestorowa [17], GSE99235 from cells in mice lung conducted by Vanlandewijck [9], and GSE107552 of human pluripotent stem cells conducted by Han [8]. All the datasets were clustered by k-NN clustering methods in Seurat after t-SNE or log transformation process into 21, 14 and 23 types correspondingly. The results were different in these three datasets (fig.1A-B). In order to analyze how the results were relevant to biology knowledge, we chose gold standard biomarkers of specific cell types in three datasets. In the dataset of Nestorowa, we used *Gata2* referring to erythrocyte-megakaryocyte precursor cells (Pre-EM) and *Elane* referring to mature granulocytes. In the dataset of Vanlandewijck, we used *Cd68* referring to macrophages and *Col2a1* referring to chondrocyte. In the dataset of Han, we used APOA1 referring to adipose lineage cells and ACTC1 referring to muscular lineage cells [8]. Given that there are many cell types in one clustering result, how each of them response to one biomarker can be measured by receiver operating characteristic (ROC). The power of clustering after transformation were evaluated by the maximum area under ROC curves (AUROC). In the dataset of Nestorowa, after log transformation, cell cluster 3 had the maximum AUROC to *Elane* while cell cluster 15 had the maximum AUROC to *Gata2*. But all of them were less than their counterparts – cell cluster 11 and cluster 17 after t-SNE transformation. In the dataset of Vanlandewijck, cell cluster 6 had the maximum AUROC to *Cd68* while cell cluster 11 had the maximum AUROC to *Col2a1*. They were also lower than their counterparts. In the dataset of Han, the maximum AUROC to APOA1 of cell clusters after the two transformation were similar but that to ACTC1 was higher after t-SNE transformation. So clustering after t-SNE transformation were more specific and sensitive to certain bio-markers (Fig.1C).

In addition, due to the difficulty of finding perfect parameters within few attempts, researchers have to adjust the parameters to generate different cluster results. It means that every possible result may have a chance to be selected to approach varied aims of researchers. Considering this uncertainty, we clustered the cells of each dataset into 12 to 32 groups and calculated their responses to given biomarkers with maximum AUROC respectively (Supp Fig. 1). Notably, in Nestorowa’s and Vanlandwijck’s, the results of maximum AUROCs of almost all clusters (*Elane, Gata2* in Nestorowa and *Cd68, Col2a1* in Vanlandewijck) increased after t-SNE transformation (Supp Fig. 1). In the dataset of Han, the maximum AUROCs of all clusters also increased or remain similar after t-SNE transformation, especially for ACTC1 (Supp Fig. 1).

**Figure 1.**
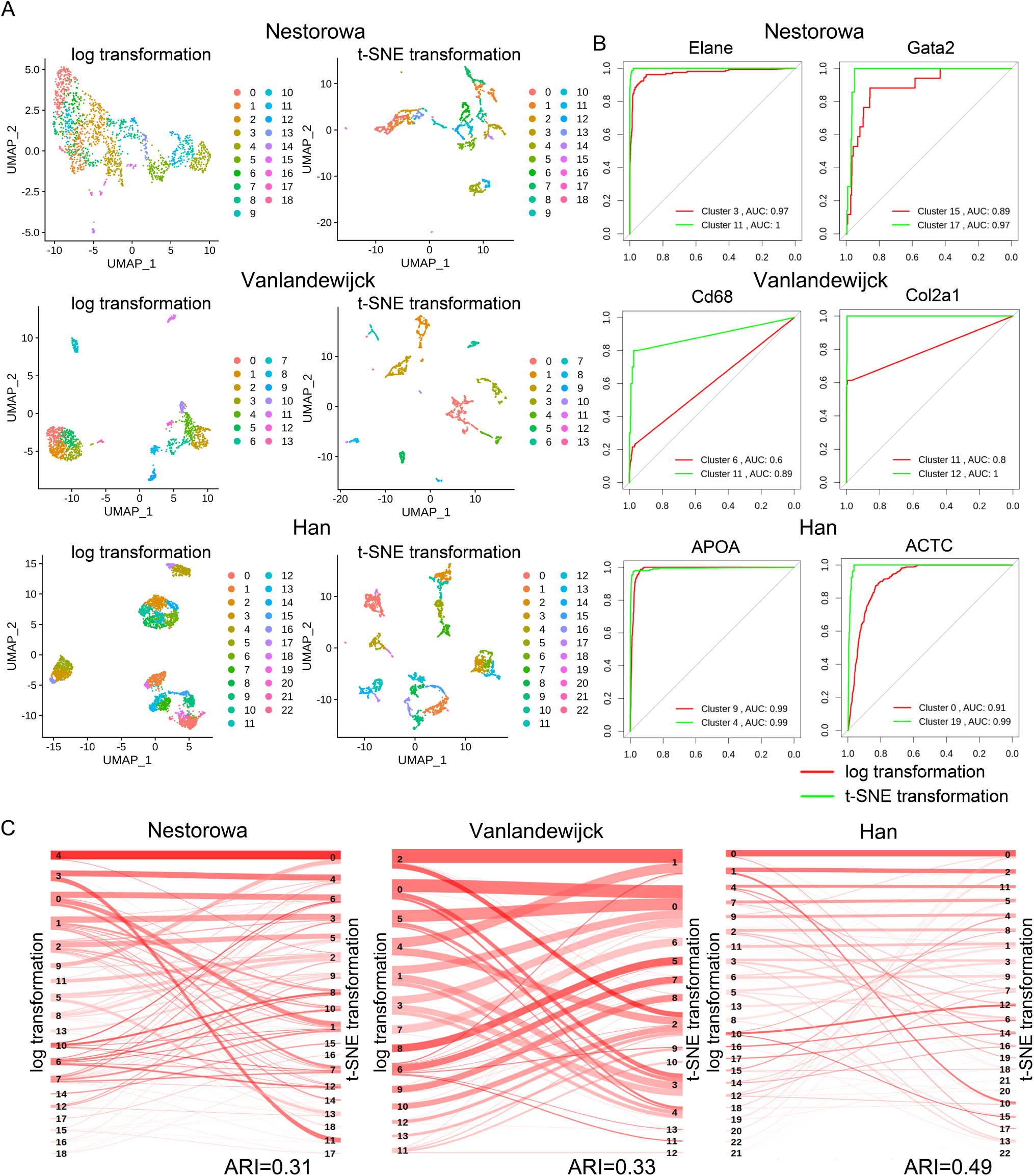
kNN clustering after log transformation or t-SNE transformation. A: Clustering results on UMAP visualization of datasets of Nestorowa, Vanlandewijck and Han after log or t-SNE transformation. B: Sankey diagram displayed different clustering results between log and t-SNE transformation. C: ROC curves demonstrated the responses of clustering to biomarker genes. Clusters with the maximum value and the maximum area under curve were labelled. The larger number of areas displayed means higher accuracy of clustering responding to biomarkers.

Therefore, these results suggested that clustering results after t-SNE transformation could infer different information from those after log transformation, and they were relevant to existed biology knowledge.

### Clustering after t-SNE transformation was steady when cluster number changed

In order to investigate the mechanisms of t-SNE transformation, we analyzed how the clustering makes the decision when the clustering number is changed after different transformation methods. This performance is named as steady of clustering in this study. It is originated from the adjusted rand index (ARI), which calculates the similarity of a clustering result to its counterpart when clustering number increase or decrease by 1. When the steady is low, it indicates that the division of a large group into balanced subgroups after clustering number is changed. Otherwise, a higher steady indicates the isolation of a small number of cells. Obviously, high steady suggested the methods being sensitive to local features of datasets. In this study, the three datasets were clustered into 12 to 32 groups described previously. The results demonstrated that the steady of clustering after t-SNE transformation were higher than those after log transformation significantly (Fig.2). Therefore, t-SNE transformation should make clustering focus on local features.

**Figure 2.**
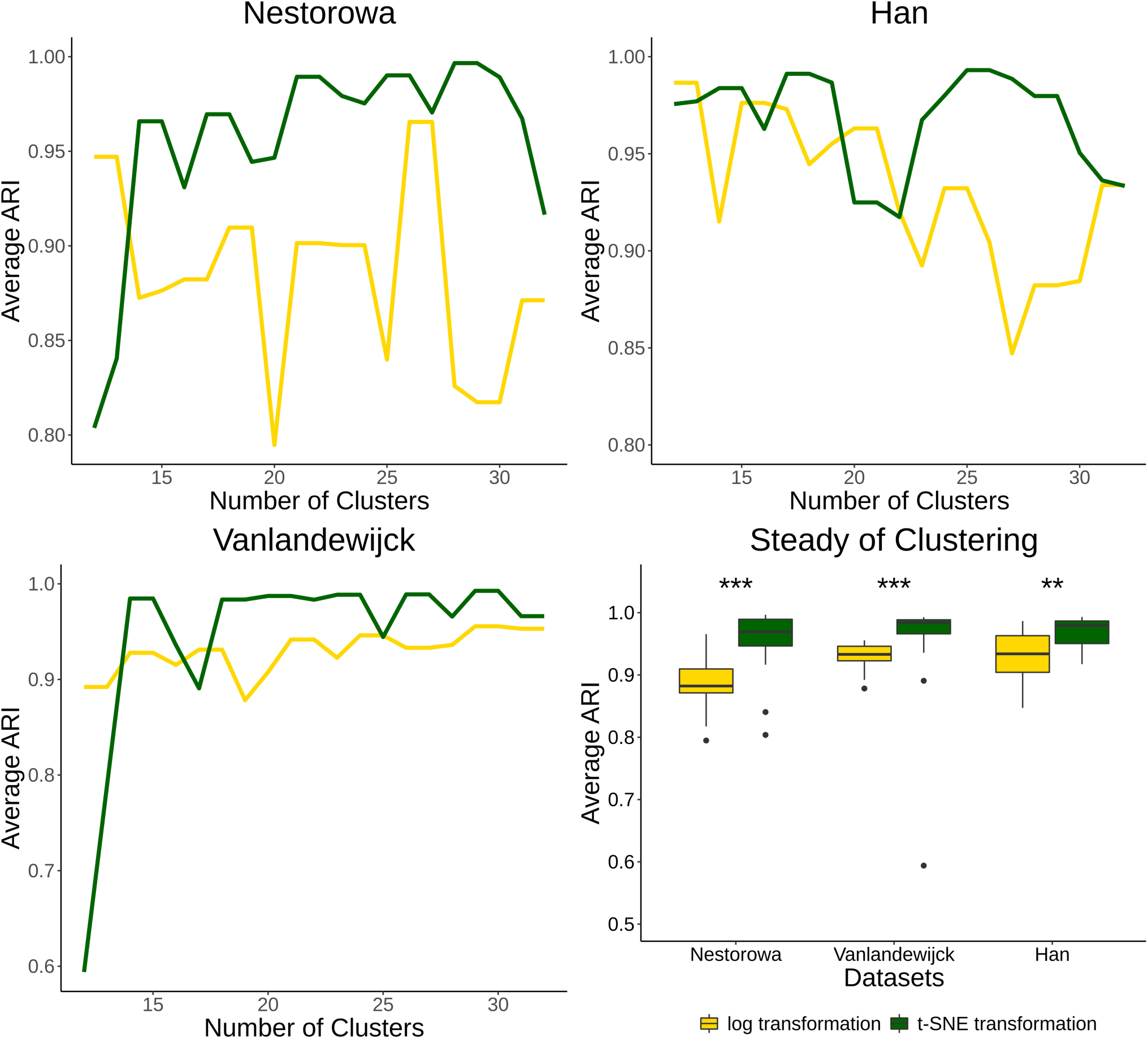
Steady of clustering with different cluster numbers. Boxplot showed the statistic results of clustering stability. Significance level: ***: adjusted p-value<0.001; **: adjusted p-value<0.01;

### t-SNE transformation was sensitive to rare cell populations

One of the characteristics of clustering oriented local features is that it is sensitive to rare cell populations. It meets the requirements of specific biological studies. With the same *k*.*para* and clustering numbers, the minimum populations of cells in clustering results after t-SNE transformation were smaller than those after log transformation (Table.1). The numbers were close to the limitation of k-NN clustering with *k*.*para* being 7. The populations of the rare cells can be interpreted biologically. For example, in the dataset of Han, there is a subpopulation with only ten cells highly expressed LAMA4, the marker of trophoblasts [19], after t-SNE transformation (Supp Fig.2). In the visualization of UMAP, they are isolated from other cells in a large scale (Supp Fig.3).

**Table 1.** Cell numbers in top 3 rarest population types

| Datasets | Order of rarity | log Transformation | t-SNE transformation |
| --- | --- | --- | --- |
| GSE81682 | 1st | 9 | 7 |
| GSE81682 | 2nd | 9 | 7 |
| GSE81682 | 3rd | 17 | 9 |
| GSE99235 | 1st | 22 | 7 |
| GSE99235 | 2nd | 32 | 7 |
| GSE99235 | 3rd | 44 | 10 |
| GSE107552 | 1st | 24 | 10 |
| GSE107552 | 2nd | 29 | 10 |
| GSE107552 | 3rd | 31 | 13 |

However, whether there were trophoblasts or any other rare cell types in the sample were not verified with the biology experiment in this study. Considering this, we removed majorities of cells in existed types in datasets of Nestorowa and Han instead. It created artificial rare cell populations with strong confidence consequently. Herein, only 15 granule-monocyte lineage cells (*Elane*+, cluster 4 and 11 after t-SNE transformation) in Nestorowa’s dataset and 15 adipose lineage cells (APOA1-high, cluster 4 after t-SNE transformation) in Han’s dataset were left for analyzing use. The UMAP showed that *Elane* and APOA1 high expressed cells aggregated and separated from other cell populations after t-SNE transformation (Fig.3A). They corresponded to group 14 in GSE81682 and group 11 GSE107552 (Supp. Fig 4). Meanwhile, they were merged into other cell types after log transformation with no cell groups’ specific expression (Fig.3A, Supp Fig.4). It was demonstrated by the larger maximum AUROC corresponding to the specific gene marker after t-SNE transformation (Fig.3B). In addition, when we changed the cluster number, clustering responded to the two markers (*Elane* or APOA1) more accurately (Fig.3C, Supp Fig 4). And the cell populations with highest expressed *Elane* or APOA1 in the corresponding dataset were significantly similar to group of primary residual cells (Fig.3D). As a result, the potential of detecting rare cell populations after t-SNE transformation is more powerful.

**Figure 3.**
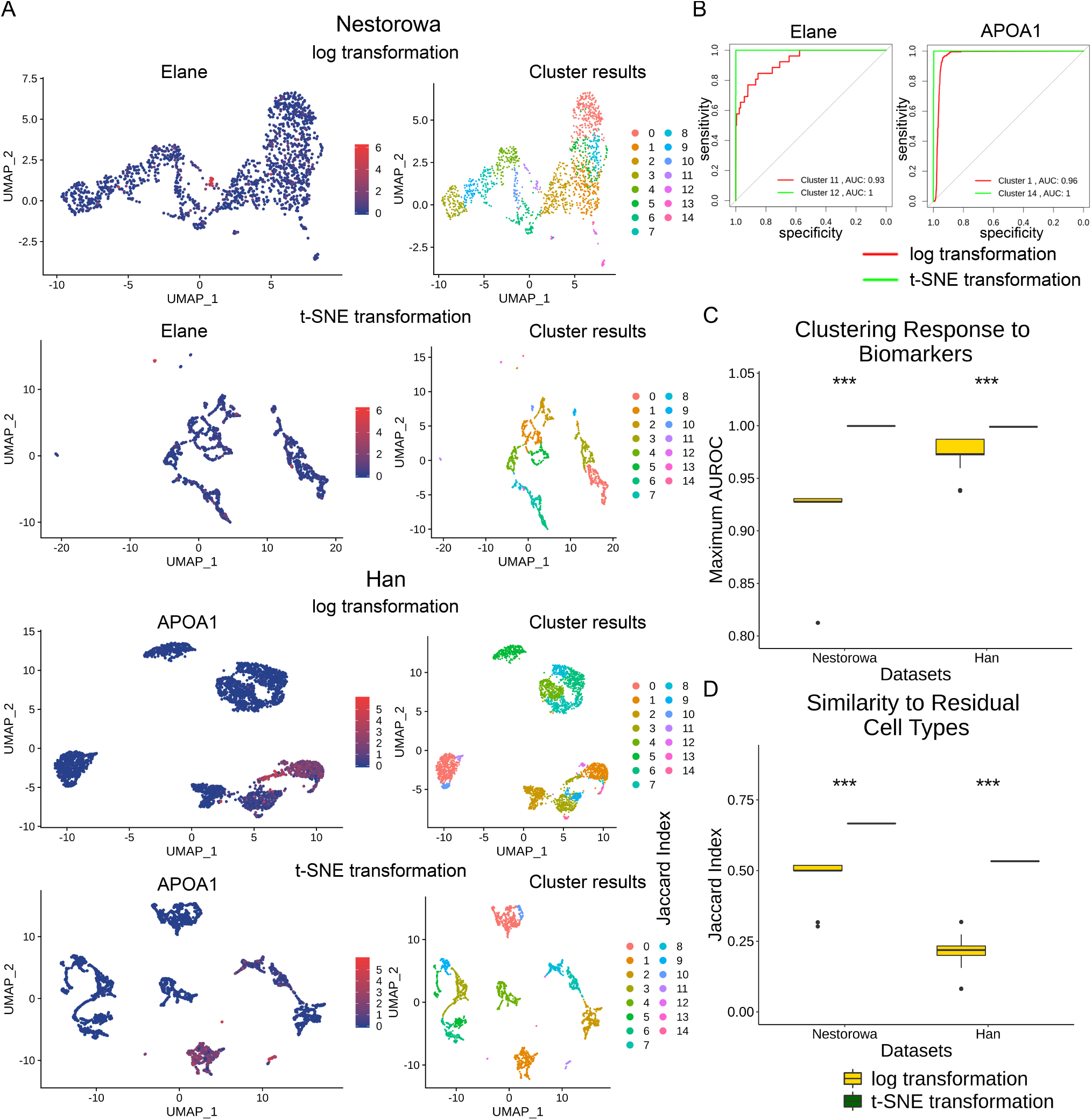
Heatmaps of differential expressed genes after delete cells. A: Heatmaps showed differential expressed genes between the cell types detected by k-NN clustering in Nestorowa after log or t-SNE transformation. B: Heatmaps showed differential expressed genes between the cell types detected by k-NN clustering in Han after log or t-SNE transformation. ***: adjusted p-value<0.001

**Figure 4.**
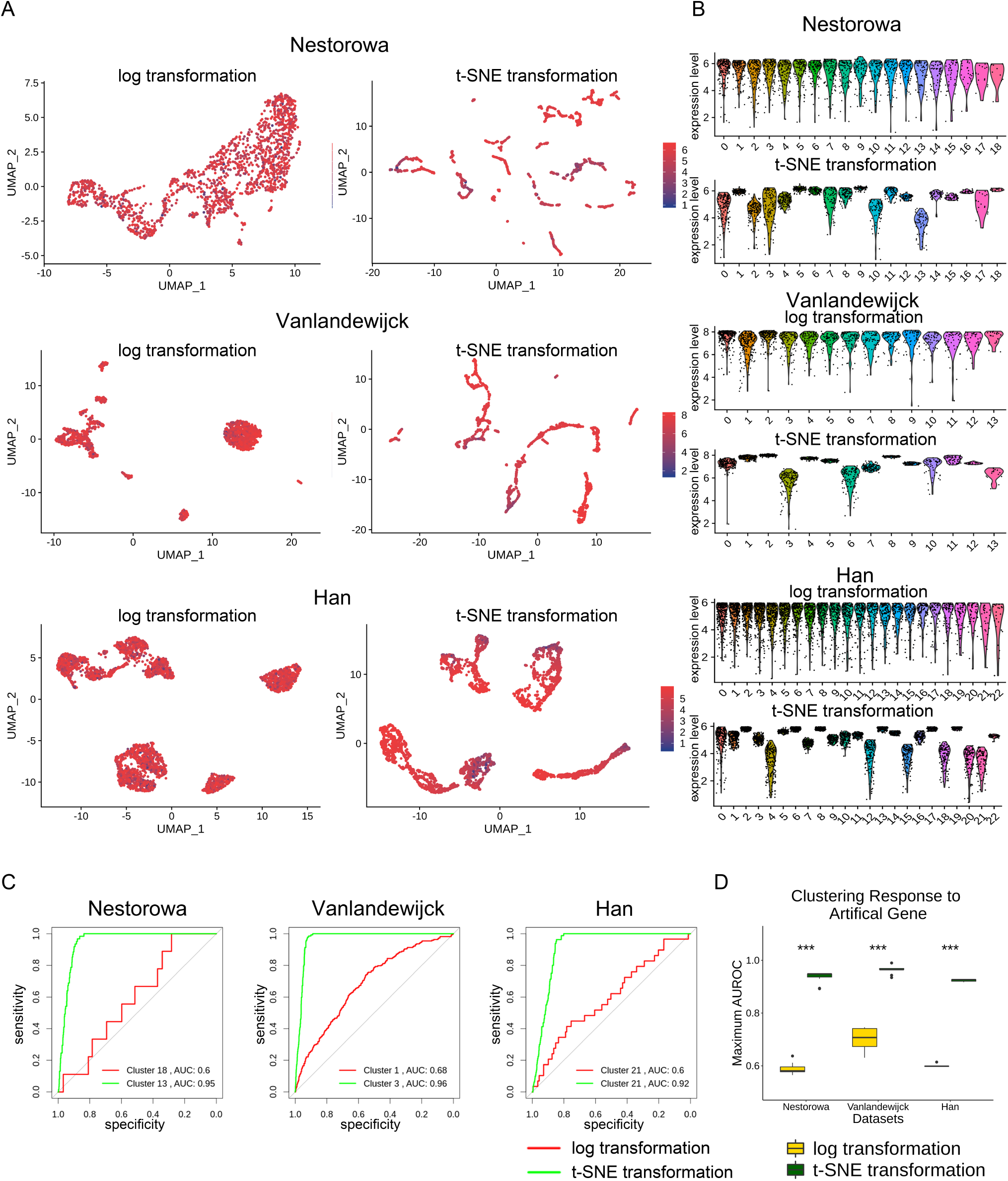
Clustering results and assessment of after log or t-SNE transformation when adding the artificial gene in datasets. A: Feature plot of artificial gene on UMAP visualization after log or t-SNE transformation in three datasets B: Violin plot showed artificial gene expression in different cell types after log or t-SNE transformation in three datasets C: ROC curves showed the responses of clustering to artificial gene. Curves were labeled by cell types with maximum area under the curve. D: Box plot showed the comparisons of clustering responses for biomarkers between log and t-SNE transformation. ***: adjusted p-value<0.001

### t-SNE transformation is sensitive to independent but highly-variated genes

The local features of datasets also mean the pattern consisted of limited genes. According to the mathematical description of t-SNE transformation, it centralizes cells based on their direct distance. t-SNE transformation will consider the variation contributed by rare genes. On the contrary, because log transformation is a concave function, variation in large scale on one gene will be reduced dramatically. In addition, if the variation of an independent gene is similar to the sum of variation of genes that correlated with each other, it will be reduced much more after log transformation.

In order to obtain a clear view, we add an artificial gene served as a relevant virtual biomarker to all datasets. It follows a uniform distribution from 0 to half of the maximum gene expression. The visualization showed cells distributed according to the expression of artificial genes after t-SNE transformation, but those after log transformation did not perform the same tendency (Fig.4A). The artificial gene also expressed differentially in different subpopulations (Fig.4B). It was shown in Heatmap of the differential expressed gene (Supp Fig.5). Sankey diagram and Heatmap also illustrate the dramatic changes of the t-SNE transformation after adding artificial gene (Supp Fig.6). Expectedly, clustering response to artificial gene was much higher after t-SNE transformation in all datasets (Fig.4C,Supp Fig.7). After log transformation, all the maximum AUROC for artificial genes were closed to 0.5, suggesting clustering results were not correlated to the artificial gene. In general, t-SNE transformation is sensitive to independent but highly variated genes.

Obviousely, in the dataset of Nestorowa, a cluster highly expressing *Gata2* was detected accurately in group 17 after t-SNE transformation, which is relavent to cell fate commitment of megakaryoerithrocyte lineage (Fig. 1B). It almost failed after log-transformation because of expression of *Gata2* may be independent to other genes describing the route of erythrocyte maturation.

## Discussion

In this study, we developed an algorithm called t-SNE transformation based on t-SNE for normalizing scRNA-seq data. We compared its performance with log transformation by using three real datasets (GSE81682, GSE99235, GSE107552) downloaded from GEO database. We found that the clustering after t-SNE transformation not only responded better to certain golden standard biomarkers, but also perform a steadier state than that after log transformation even when the clustering parameters were changed. In addition, the t-SNE transformation made detection of residual cells more accurate after deleting the majority of specific cell types from the original datasets. Further study also showed that when adding an artificial gene that expressed independently to the real genes, the t-SNE transformation was sensitive to it while log transformation was not.

As described above, different scRNA-seq data analysis tools are applied to meet diversified research objectives. It will benefit the whole field of scRNA-seq study if algorithm designers can provide specific tools targeted certain research purposes. Not only its evaluation level would be judged by the statistic assessment criterion, but also its preferential target dataset should be clarified. In this study, t-SNE transformation is designed for aims to local features: cell types with rare population and highly variated independent features.

Since the mechanisms of t-SNE is an analogy of sigmoid normalization in supervised machine learning, samples can aggregate to different cluster centers after t-SNE transformation. It means that if a small group of cells is far away from the general population, they will aggregate with each other and form a more detectable cluster center than those without t-SNE transformation. That is also the reason why the isolation of small subpopulations is prior to the division of a large population cell group after t-SNE transformation. When the clustering number increased, the results become steadier. Expectedly, we detected 10 Cd68 positive cells in GSE99235 and 10 LAMA4 positive cells in GSE107552 after t-SNE transformation while log transformation failed to detect them. We also detected the artificial rare population cell types which were generated by removing most of cells in one of main types in datasets. According to a tool developed by Christoph Hafemeister https://satijalab.org/howmanycells based on negative binomial distribution, the possibility of detection should be less than 5

Besides, cell types in datasets may be different from each other semantically, which means that each of them can be marked by numerous genes. If they are relevant to the aims of studies, log transformation will work well since it overestimates the sematic variation of transcriptome. It still cannot meet the requirements of certain biological studies such as discovering the new functional cell types or critical regulation genes. Supposed that the artificial gene generated in this study is an essential candidate for particular biology processes, it may be ignored when applying the log transformation. In addition, Cd68 was high variated between blood and non-blood cells in the dataset of Vanlandewijck but was underestimated by log transformation. Considering this, we suggest that t-SNE transformation can be a solution to solve this problem.

It is the characteristics of t-SNE transformation that are sensitive to local features rather than the advantage. It is not only decided by various aims of biology studies, but also by considering the high noise in scRNA-seq datasets. The noise may be generated from the library construction process or during the sequencing procedure.

It may also generate highly variated independent genes or the cell types with rare populations and cause pseudo positive results. In this situation, the efficiency and accuracy of t-SNE transformation could be undermined while log transformation should be better than t-SNE transformation. because of this, t-SNE transformation can be used in a prior study of a project rather than a systemic summary. Its results should also be verified by other biological experiments.

In conclusion, t-SNE transformation is a normalization method for local features. It is suitable for the study dedicated to finding rare population cell types or highly variated but independent genes. Log transformation could not provide enough information under this condition.

## Methods

Data quality control, preprocessing and log normalization were discribed in supplementary methods

### t-SNE transformation

As its name, t-SNE transformation was derived from t-SNE [16]. In detail, the mathematic process can be described below:

The possibility of linkage between cell i (*x*_*i*_) and cell j (*x*_*j*_) is defined as:

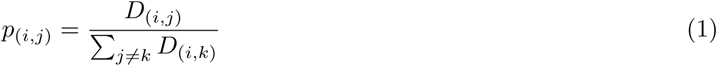

where *D*_(*i,j*)_ is the distance between cell i and cell j before transformation.

Similarly, the possibility between cells after t-SNE transformation is defined as:

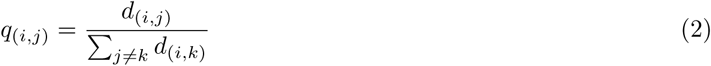

where *d*_(*i,j*)_ is the distance between cell i and cell j after transformation.

In t-SNE, D is in high dimension, while d is in low dimensions such as 2-D or 3-D for dimension reduction or visualisation [16]. In this study, t-SNE transformation changes the dimension of d equal to that of D. It is the main difference from t-SNE to t-SNE transformation.

Because the denominators are different in different cells, to simplify this, they can be replaced by the total sum of distances between all samples.

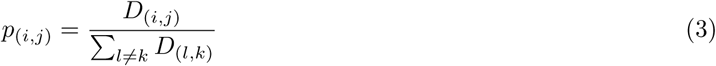

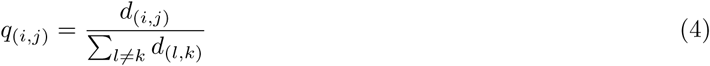

Let ***p***≃***q*** and sum of KL divergence of all vectors in ***p*** from corresponding vectors in ***q*** is used as cost function:

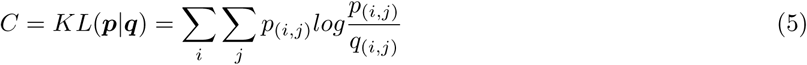

Then the Gaussian kernel is introduced to *D*_(*i,j*)_:

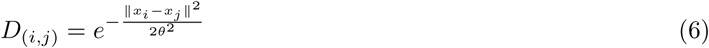

where *x*_*i*_ is genes expression vector of cell i

And the Cauchy kernel is introduced to *d*_(*i,j*)_:

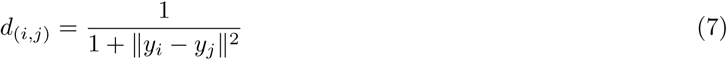

where *y*_*i*_ is the location of cell i after t-SNE transformation.

In this situation, the cost function for Gradient descending become:

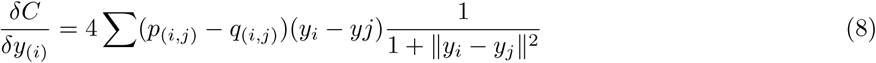

In gradient descending, ***y***_**(0)**_ is initialised randomly. Then:

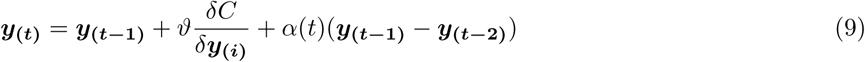

where *θ* is learning rate and *α*(*t*) is a function of t to make *C* does not fall into the local minimum.

According to those descriptions, libraries of t-SNE transformation are the same as t-SNE written by Maarten [16]. Considering the cost of time, we ran it in MATLAB 2017a. Its codes were downloaded from https://lvdmaaten.github.io/tsne/. Unlike dimensionality reduction with t-SNE, ‘no_dim’ (final dimension) in the function was set the same as ‘initial_dim’ (initial dimension). Then the results were written to csv file for further analysis.

### Comparing clustering performance

#### Clustering response to biomarkers

To investigate whether clustering was relevant to some exist or potential biological knowledge, we chose some standard gold biomarkers as indicators and test whether they matched one of the cell groups in clustering results. In Nestorowa, they were Gata2, the marker of Megakaryocyte–erythroid progenitor cells (pre-ME) and Elane, the marker of Granulocytes. In Vanlandewijck, they were Cd68 the marker of dust cells and Col2a1, the marker of chondrocytes. In Han, they were APOA1, the marker of adipose lineage cells and ACTC1, the marker of muscular lineage cells [8]. In supervised clustering, receiver operating characteristic (ROC) can be used as an evaluation of classification with features. The more area under ROC curve (AUROC), the more correct classification response to the feature [24]. Similarly, it can also be used to evaluate how the clustering responses to specific markers. Herein, it was defined as the maximum AUROC in all cell types detected by clustering. All the cluster results were assessed by maximum AUROC on biomarkers above under the clustering conditions mentioned previously.

Furthermore, because users are not certain of the best clustering parameters in their own studies, investigation of clustering performance should consider more parameters. So we divided cells in these three datasets into 12 to 32 groups after log or t-SNE transformation to involve more possible clustering results. The responses of all clustering results to corresponding biomarkers listed above were calculated with the maximum AUROC and shown in boxplot.

#### Steady of clustering

Given the cluster number is the essential parameter of clustering, we defined the steady of clustering as:

Considering A is a clustering result, steady of clustering of A refers the average of its similarities to the clustering results whose cluster number were one more and less than that of A. In this study, the clustering similarity is defined as the adjusted rand index (ARI) [13]:

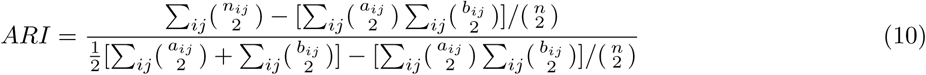

where *i* and *j* represent sets of cells in clustering result with different method or parameter, *n*_*ij*_ = *|i ∩ j|, a*_*i*_ =∑ _*j*_ *n*_*ij*_, *b*_*j*_ = _*in*_*ij*_, and 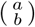 means the binomial coefficient.

In this study, this performance was assessed when cells in datasets were divided into 12 to 32 groups. The result with every cluster number in one dataset can generate one steady of clustering. All the results were displayed with line-plot and boxplot.

#### Sensitivity to the rare population

The rare cell populations were created by deleting majorities of one cell type with concrete function in datasets of Nestorowa and Han randomly. They had been detected by clustering after both transformations with original datasets. Specifically, they are Granulo-monocytes lineage cells in the dataset of Nestorowa and Apidose lineage cells in the dataset of Han to 15. Both of them occupied over 10% population in the corresponding dataset. After that, we used the same methods and parameters described above and tested how clustering response to Elane or APOA1, the markers of corresponding cell lineages, with ROC. We also tested this characteristic of clustering when cluster numbers were set from 12 to 32. As well as comparing the response of every clustering with different cluster number to Elane or APOA1 correspondingly, the similarities between cell types with max AUROC of these two markers to corresponding residue cells were calculated with the Jaccard Index:

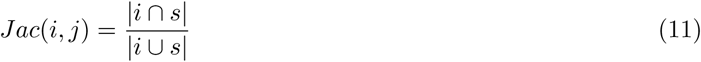

where *i* is cell type with the maximum AUROC to the corresponding biomarker (*Elane* or APOA1) in clustering results and *s* is residual cells.

All the results were shown with boxplot.

#### Sensitivity to high variated but independent gene

An artificial gene was added to the original datasets and evaluated how they affected clustering results after log or t-SNE transformation. The artificial gene followed unique distribution and expressed randomly in cells. After that, we used the same transformation and clustering methods described above. Then we tested how clustering response to this artificial gene with ROC under the conditions with cluster number set from 12 to 32. All the maximum AUROC of every clustering results were shown in boxplot.

#### Statistics

All the significance of difference was calculated with Kruskal-Walli test. Pair comparisons used Dunnet’s test. P-value was adjusted by the false discovery rate (FDR). The significance level was 0.05.

## Supporting information

Supplementary methods and results

## Additional information

All authors do not have competing interests.

