## Supplementary methods and results for "t-SNE transformation: a normalization method for local features of single-cell RNA-seq data"

### Datasets and Quality Control

Datasets were downloaded from the GEO database (GSE81682, GSE99235, and GSE107552. Quality control was applied to remove errors, eliminate irrelevant information as well as to avoid the effect of low-quality samples. This process was run by MATLAB R2017a. Genes that were not annotated with “protein_coding” in Ensembl were removed. Ribosomal protein genes were also excluded because their variation did not provide enough relevant information to recognize cell types. Cells with too high or low count numbers or features numbers were removed.

In datasets of Nestorowa and Vanlandewijck, researchers used Smart-Seq2 to construct libraries [2, 3]. It fragmented the whole mRNA before amplification [4]. As a result the transcript numbers of mRNA is correlated to its length. Herein, we used the Transcripts per million base (TPM) to calculate the expression matrix:

$$TPM_{(i,j)} = \frac{RPKM_{(i,j)}}{\sum_i RPKM_{(i,j)}} = \frac{\frac{C_{(i,j)}}{L_i}}{\sum_i \frac{C_{(i,j)}}{L_i}} \tag{1}$$

where RPKM means Reads Per Kilobase Million, $i$ is the identity of genes, $j$ is the identity of cells, $C_((i,j))$ is the count number of fragments mapping to gene $i$ in cell $j$ and $L$ is the total length of exomes of gene $i$.

In datasets of Han, researchers used the C1-800 platform. This method used 3’ fragments to construct libraries. It means one read represents one transcript. The length of mRNA should not be involved in calculation. Therefore, we used the Counts of exon model per million mapped reads (CPM) to calculate the expression matrix:

$$TPM_{(i,j)} = \frac{C_{(i,j)}}{\sum_i C_{(i,j)}} \tag{2}$$

where $i$ is the identity of genes, $j$ is the identity of cells,$C_((i,j))$ is the count number of fragments mapping to gene $i$ in cell $j$.

Finally, genes with CPM or TPM largern than two in at least two cells were kept in this study.

### Pre-processing, Visualization, Clustering, and Heatmap in Seurat

Seurat is one of the most popular scRNA-seq data analysis platform with a friendly interface for external programs [5]. It involves log transformation procedure which can act as the counterpart of t-SNE transformation. Clustering and visualization are also implemented in Seurat.They can be launched easily after either transformation method. So we used the Seurat for continuing analysis based on its advantages.

The preparation processes followed the instruction of Seurat in general. In detail, after loading the expression matrix, a Seurat object was created, then the data was normalized by setting the method to "LogNormalize" while *scale.factor* to 10000. The feature selection process is skipped to avoid repetitive operation. We ran "ScaleData", "RunPCA" and "JackStraw" functions with the default setting, which are mandatory/default for following steps. When the object was copied, we replaced the dimension reduction table in the copy with csv files of t-SNE transformation results.

Datasets after t-SNE or log transformation underwent the same clustering and visualization pipeline, then they were clustered with k-NN clustering method. The k.params were set to 7 to increase the sensitivity to rare cell types. Low dimension visualization was completed with UMAP in Seurat [1]. The differences between clustering results after log and the t-SNE transformation were visualized with Sankey diagram.

In order to obtain a clear understanding of the biological meaning of cell groups in the clustering results, we selected the differentially expressed genes in each type of cells with "FindAllMarkers" function of Seurat. Wilcoxon Ranking Sum Test was chosen to accomplish the task. The p-value threshold was set to 0.05. The logFc threshold was set to 0.25. Heatmap was illustrated by the "DoHeatMap" function. It used the top four log fold changes among differentially expressed genes in each group.

### Supplementary Figures

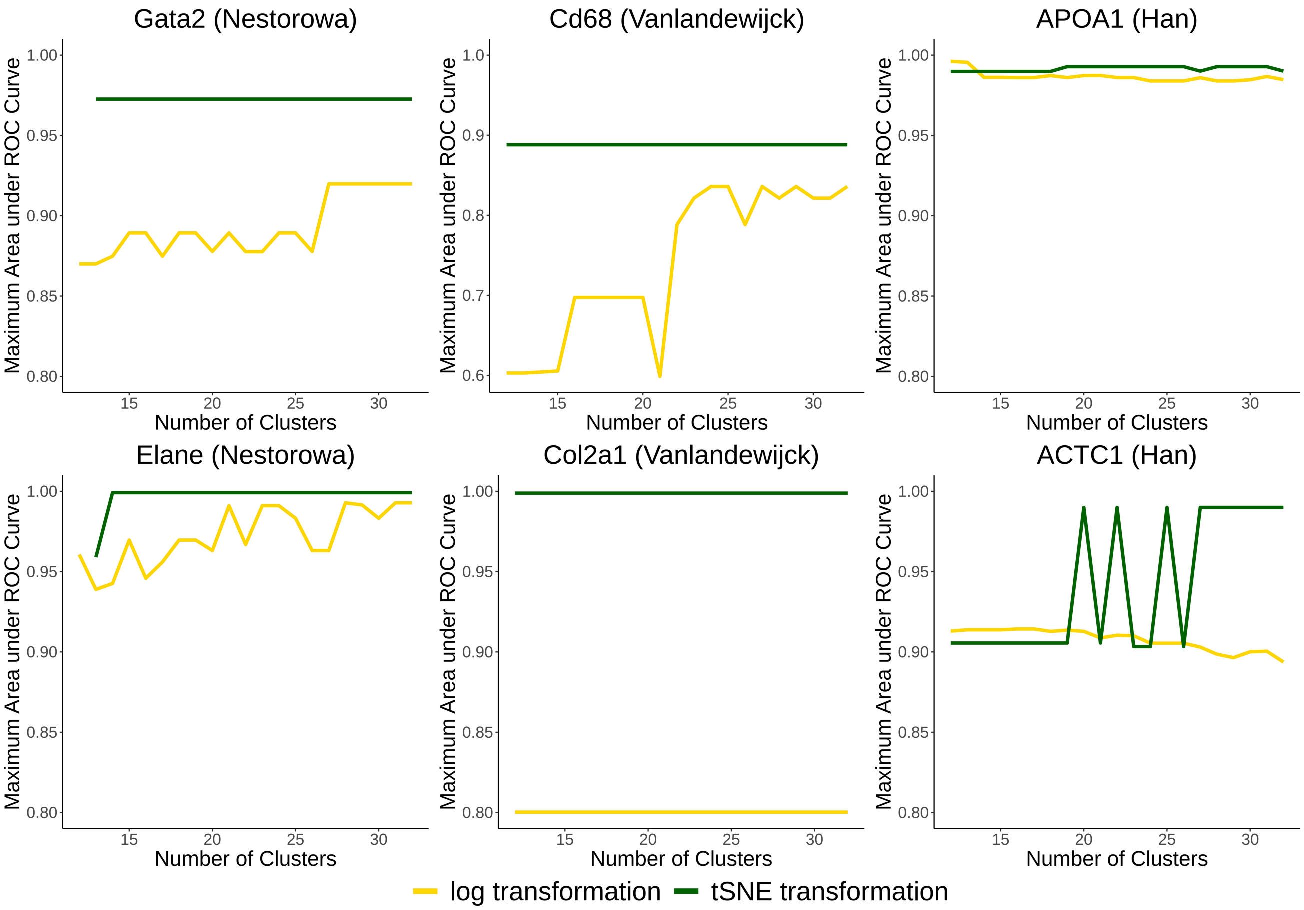


**Supplementary Figure 1: Clustering responses to corresponding markers with different cluster numbers.**

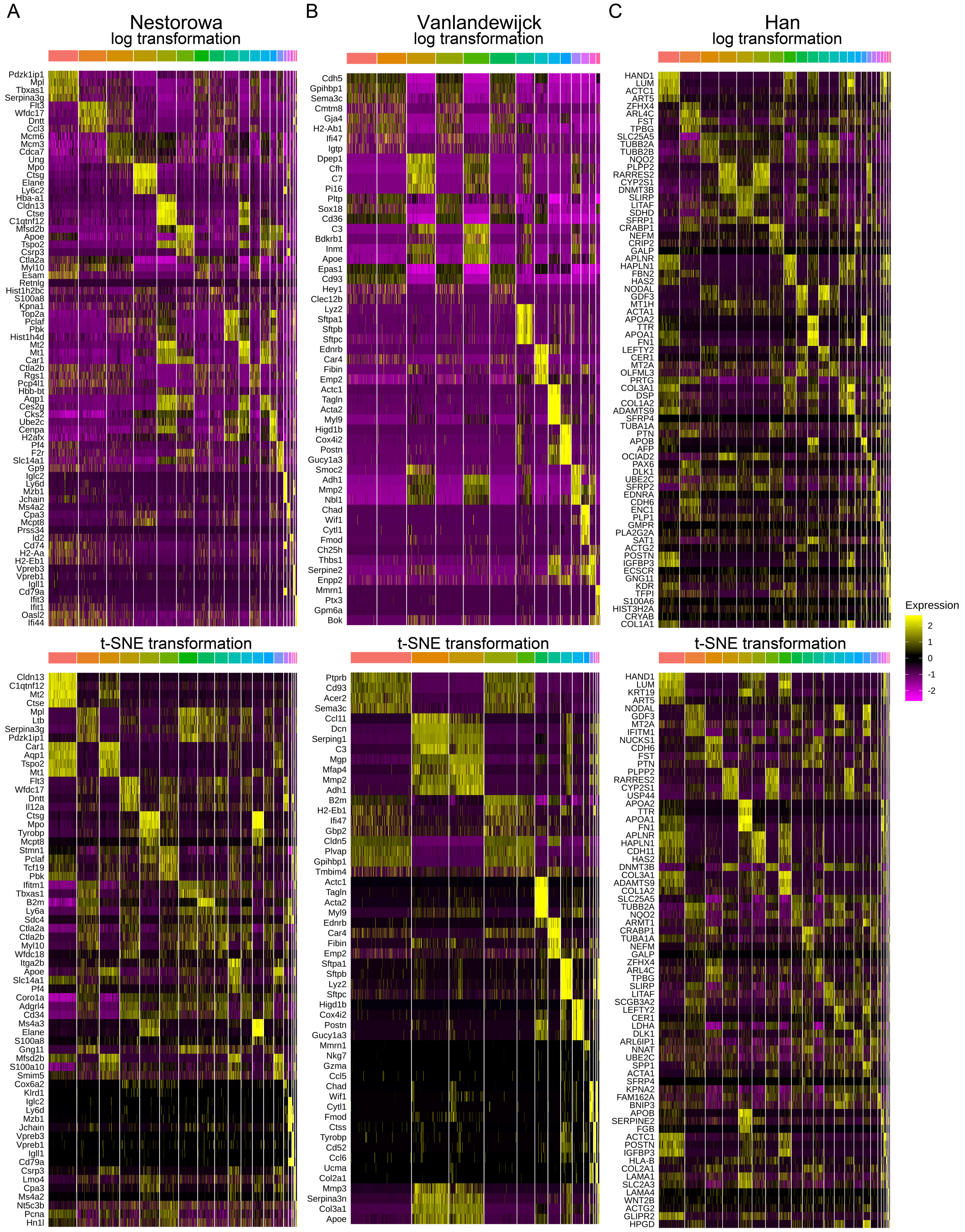


**Supplementary Figure 2: Heatmaps of differential expressed genes.** A: Heatmaps showed differential expressed genes between cell types detected by k-NN clustering in Nestorowa after log or t-SNE transformation. B: Heatmaps showed differential expressed genes between cell types detected by k-NN clustering in Vanlandewijck after log or t-SNE transformation.C: Heatmaps showed differential expressed genes between cell types detected by k-NN clustering in Han after log or t-SNE transformation.

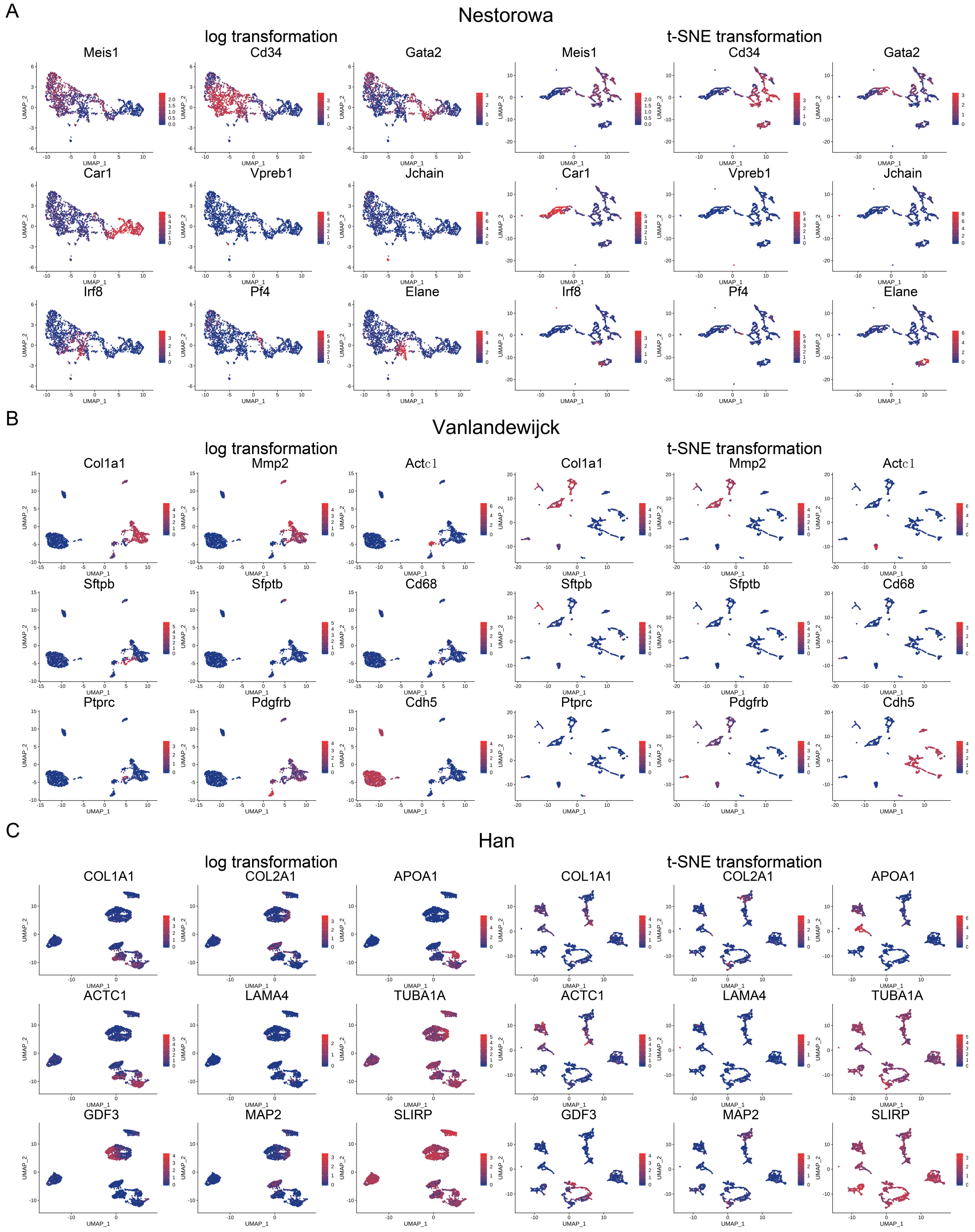


**Supplementary Figure 3: Feature scatter plots of biomarkers corresponding to datasets.** A: Feature plots of UMAP showed expression of biomarkers on cells after log transformation or t-SNE transformation of Nestorowa. Meis1: Early HSC marker; Cd34: Hematopoietic stem cell marker; Gata2: Megakaryo-Erythrocyte precursor marker; Car1: immature erythrocyte marker; Vpreb1: immature B cell marker; Jchain: pre-immune B cell marker; Irf8: monocyte marker; H2-Aa: mature monocyte marker; Elane: mature granulocyte marker. B: Feature plots of UMAP showed expression of biomarkers on cells after log transformation or t-SNE transformation of Vanlandewijck. Col1a1: Alveolar Type 1 cells (Extracellular Synthesis Marker). Mmp2: Alveolar Type 1 cells (Extracellular Degradation Marker). Actc1: Muscular cell marker. Sftpb: Alveolar Type 2 cells marker. Chad: Condrocytes marker. Cd68: Dust cell marker. Ptprc: peripheral blood cell marker. Pdgfrb: pericytes markers. Cdh5: Blood vessel cells marker C: Feature plots of UMAP showed expression of biomarkers on cells after log transformation or t-SNE transformation of Han. COL1A1: connective tissue markers. COL2A1: Chondrocyte lineage marker. APOA1: Adipose lineage cell marker. ACTC1: Muscle lineage cell marker. LAMA4: Potential trophoblast Markers. TUBA1A: cytoskeleton marker. MAP2: Neural cell lineage marker.

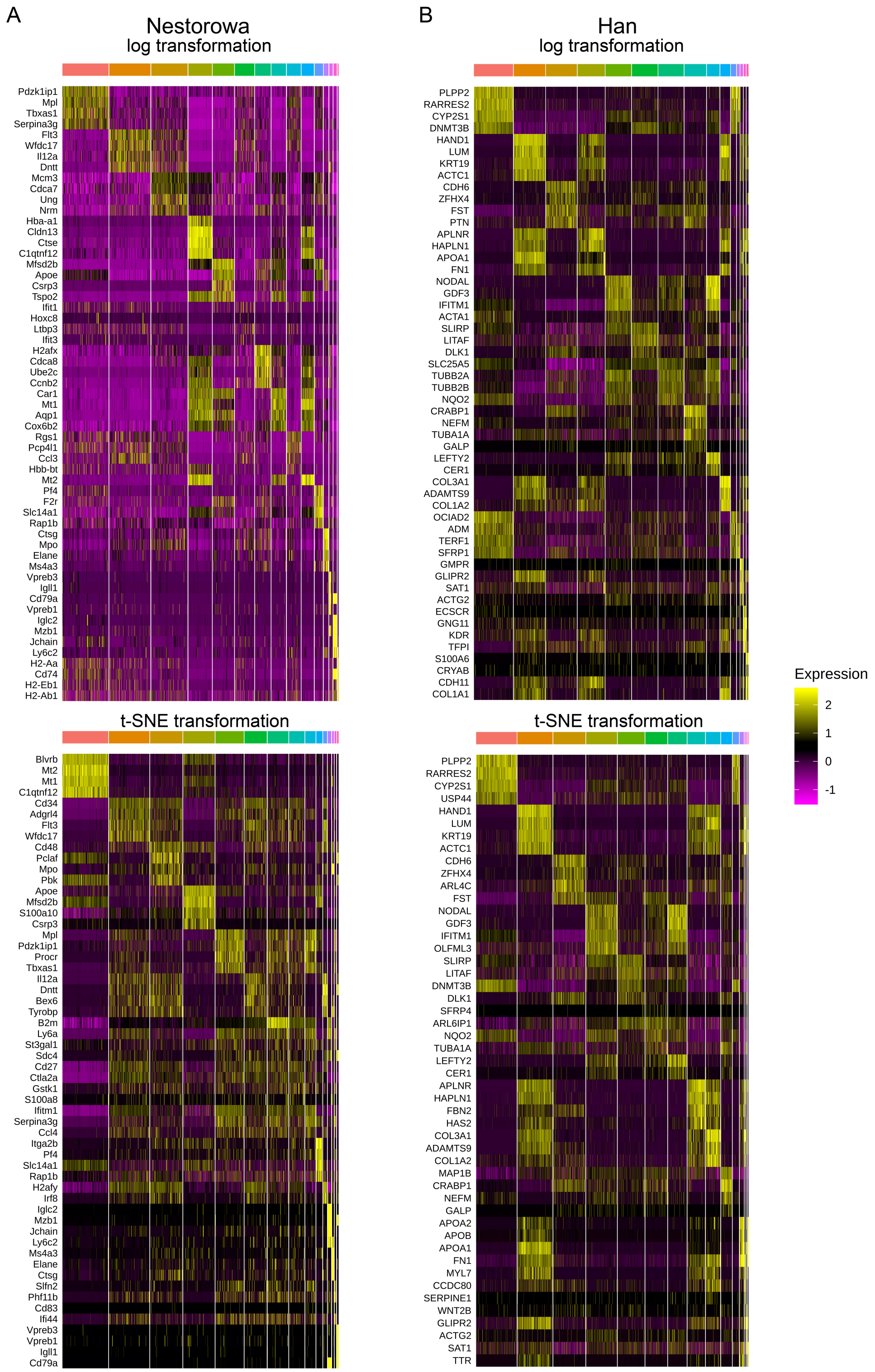


**Supplementary Figure 4: Heatmaps of differential expressed genes after delete cells.** A: Heatmaps showed differential expressed genes between cell types detected by k-NN clustering in GSE81682 after log or t-SNE transformation. B: Heatmaps showed differential expressed genes between cell types detected by k-NN clustering in GSE107552 after log or t-SNE transformation.

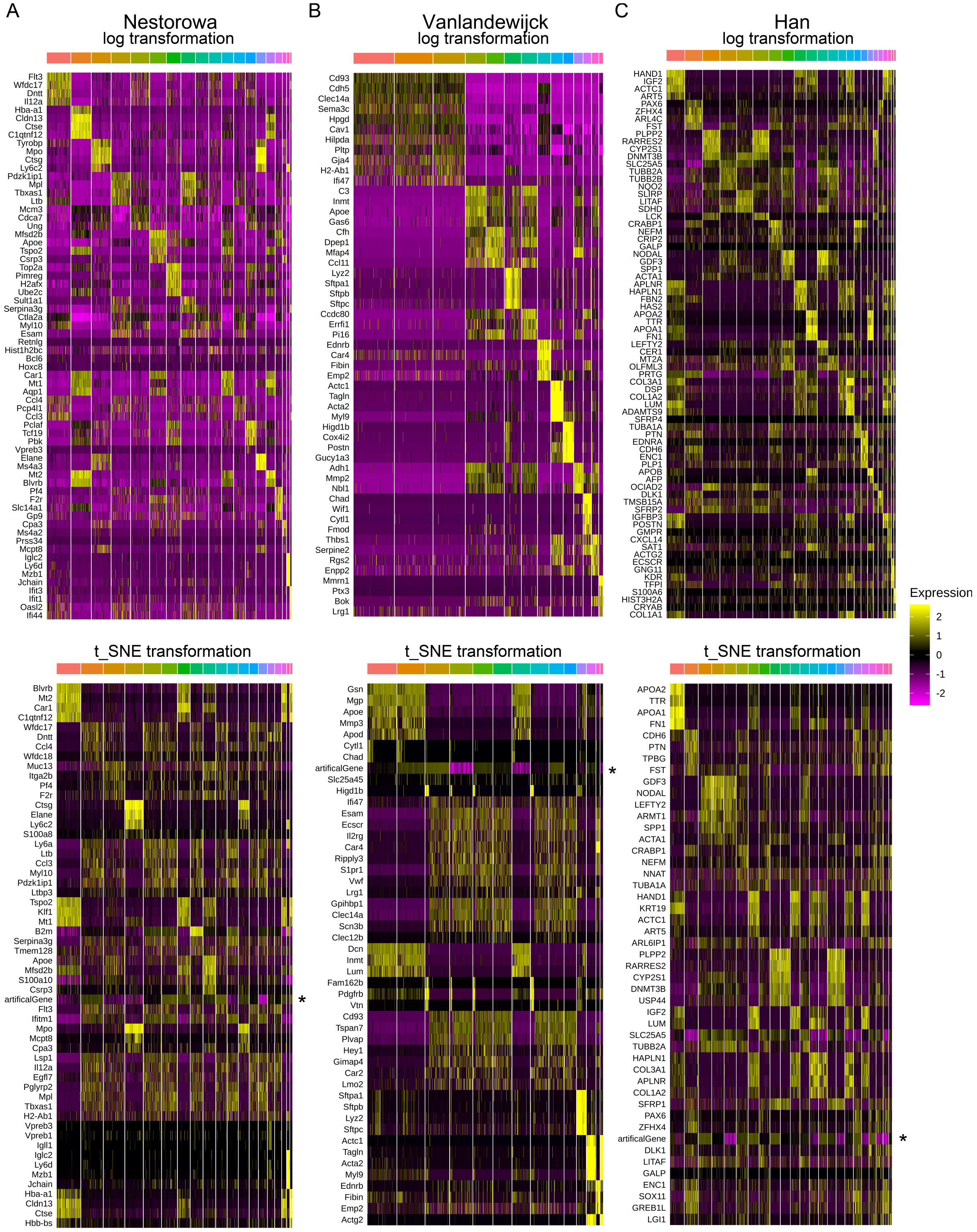


**Supplementary Figure 5: Heatmaps of differential expressed genes after adding the artifical gene.** A: Heatmaps showed differential expressed genes between cell types detected by k-NN clustering in Nestorowa after log or t-SNE transformation.B: Heatmaps showed differential expressed genes between cell types detected by k-NN clustering in Vanlandewijck after log or t-SNE transformation. C: Heatmaps showed differential expressed genes between cell types detected by k-NN clustering in Han after log or t-SNE transformation. *: location of the artificial gene in the heatmap.

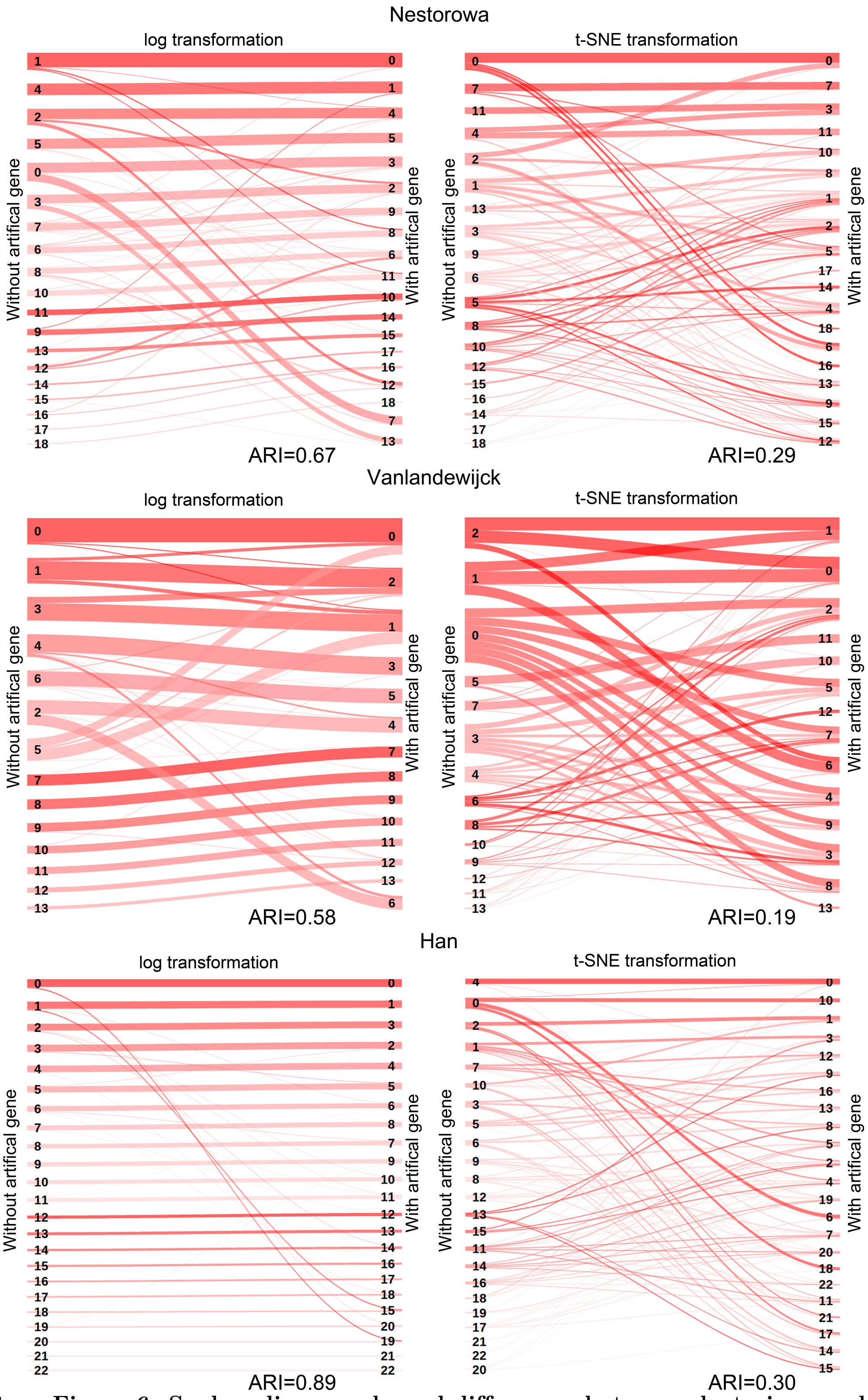


**Supplementary Figure 6: Sankey diagram showed differences between clustering results with and without the artificial gene.**

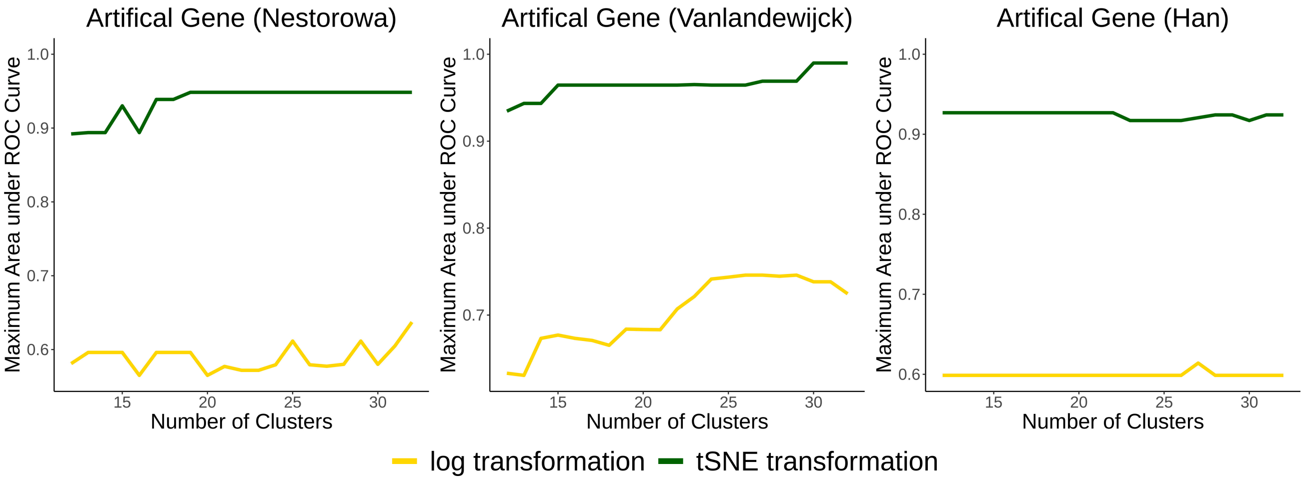


**Supplementary Figure 7: Clustering responses to the artificial gene with different cluster numbers.**
